# Mapping Antarctic suspension feeder abundances and seafloor food-availability, and modelling their change after a major glacier calving

**DOI:** 10.1101/333955

**Authors:** Jan Jansen, Nicole A. Hill, Piers K. Dunstan, Eva A. Cougnon, Benjamin K. Galton-Fenzi, Craig R. Johnson

## Abstract

Seafloor communities are a critical part of the unique and diverse Antarctic marine life. Processes at the ocean-surface can strongly influence the diversity and abundance of these communities, even when they live at hundreds of meters water depth. However, even though we understand the importance of this link, there are so far no quantitative spatial predictions on how seafloor communities will respond to changing conditions at the ocean surface.

Here, we map patterns in abundance of important habitat-forming suspension feeders on the seafloor in East Antarctica, and predict how these patterns change after a major disturbance in the icescape, caused by the calving of the Mertz Glacier Tongue. We use a purpose-built ocean model for the time-period before and after the calving of the Mertz-Glacier Tongue in 2010, data from satellites and a validated food-availability model to estimate changes in horizontal flux of food since the glacier calving. We then predict the post-calving distribution of suspension feeder abundances using the established relationships with the environmental variables, and changes in horizontal flux of food.

Our results indicate strong increases in suspension feeder abundances close to the glacier calving site, fueled by increased food supply, while the remainder of the region maintains similar suspension feeder abundances despite a slight decrease in total food supply. The oceanographic setting of the entire region changes, with a shorter ice-free season, altered seafloor currents and changes in food-availability.

Our study provides important insight into the flow-on effects of a changing icescape on seafloor habitat and fauna in polar environments. Understanding these connections is important in the context of current and future effects of climate change, and the mapped predictions of the seafloor fauna as presented for the study region can be used as a decision-tool for planning potential marine protected areas, and for focusing future sampling and monitoring initiatives.

## 1 Introduction

Primary productivity is at the base of most marine ecosystems. In Antarctica, primary production is highly seasonal and intricately tied to the location, timing and duration of sea-ice and ice-free areas such as polynyas (Arrigo and van Dijken, 2003). The collapse of large ice-shelves or calving of massive icebergs, and the retreat of sea-ice that is mainly observed around the Western Antarctic Peninsula in recent years (Parkinson and Cavalieri, 2012), can dramatically alter the oceanographic setting with down-stream effects on the pattern of primary production hotspots and on Southern Ocean ecosystems (Arrigo et al., 2002; Gutt et al., 2011b). Resulting changes in the location, timing and intensity of phytoplankton blooms (Cape et al., 2014) can influence the distribution of krill and predator aggregations (Gutt et al., 2011a), the abundances of benthic suspension feeders (Fillinger et al., 2013; Gutt et al., 2013) and can affect carbon storage (Peck et al., 2010).

For most seafloor communities living below the photic zone (∼200 m), surface-derived primary production represents their main food source (Dayton and Oliver, 1977; Duineveld et al., 2004; Ruhl et al., 2014), and is therefore critical for their survival. Seafloor communities represent the richest component of Antarctic biodiversity (Griffiths, 2010), are highly endemic (Griffiths et al., 2009), and play an important role in the marine ecosystem (Thurber et al., 2014). However, despite evidence that a changing environment influences the distribution of these communities (Gutt et al., 2011a; Fillinger et al., 2013; Gutt et al., 2013; Griffiths et al., 2017), no study has so far quantified and mapped how their distribution might change due to a changing icescape at the ocean surface. One of the reasons for the lack of quantitative studies is that although surface-derived food is one of the main drivers, it is only recently that the nature and strength of this relationship has been quantified on the Antarctic shelf using a so-called Food-Availability-Model (FAM)(Jansen et al., 2018). Combining surface-productivity and ocean currents with particle-tracking, FAMs estimate the distribution of surface-derived food at the seafloor, and evaluate the estimates against data from sediment cores. Jansen et al.(2018) demonstrated a strong link between modelled flux of suspended food along the seafloor and abundances of sessile suspension feeders, providing a framework that allows to estimate the distribution of key elements of the seafloor community and to predict how they may change with changing ocean productivity and currents.

One Antarctic region that has recently undergone drastic environmental changes is the George V shelf in East Antarctica. The calving of the Mertz Glacier Tongue (MGT) in 2010(Young et al., 2010) has resulted in profound environmental changes in the region, such as increased sea-ice concentrations (Campagne et al., 2015), and changes in ocean currents along the shelf as suggested by observations (Aoki et al., 2017) and modeling studies (Cougnon et al., 2017; Kusahara et al., 2017). These environmental changes have consequences for the dynamics and distribution of primary production (Shadwick et al., 2013), the abundance of top-predators (Wilson et al., 2016), and has been observed to influence the community structure of shallow-water benthos (Clark et al., 2015). However, the effect of the MGT-calving on the seafloor across the region has so far neither been assessed nor observed, and so its impact on benthic communities across the continental shelf is still unknown. Obtaining this knowledge, however, is crucial for meaningful assessment of the comprehensiveness, effectiveness and representativeness of the proposed marine protected areas in this region.

Here, we (i) quantify differences in the environmental setting on the George V shelf that will affect the supply of food to the benthos. In our modelling, we apply a recently developed FAM(Jansen et al., 2018) on two 5-year climatologies of remotely sensed surface chlorophyll-a for the period before and after the glacier calving, and use ocean current velocities from a purpose-built oceanographic model (Cougnon et al., 2017) (more details can be found in the Methods). We then (ii) map the distribution of benthic suspension feeder abundances before the glacier calving, using faunal abundances derived from underwater camera images and environmental predictor variables. Using the pre-calving statistical model for the suspension feeder abundances and the change in environmental conditions after the glacier calving, we then (iii) predict changes in suspension feeder abundances across the region, revealing the strong impact of the changing icescape on the seafloor ecosystem (Fig. 1 for general results, and Fig. 2 for an overview of the region).

**Figure 1.**
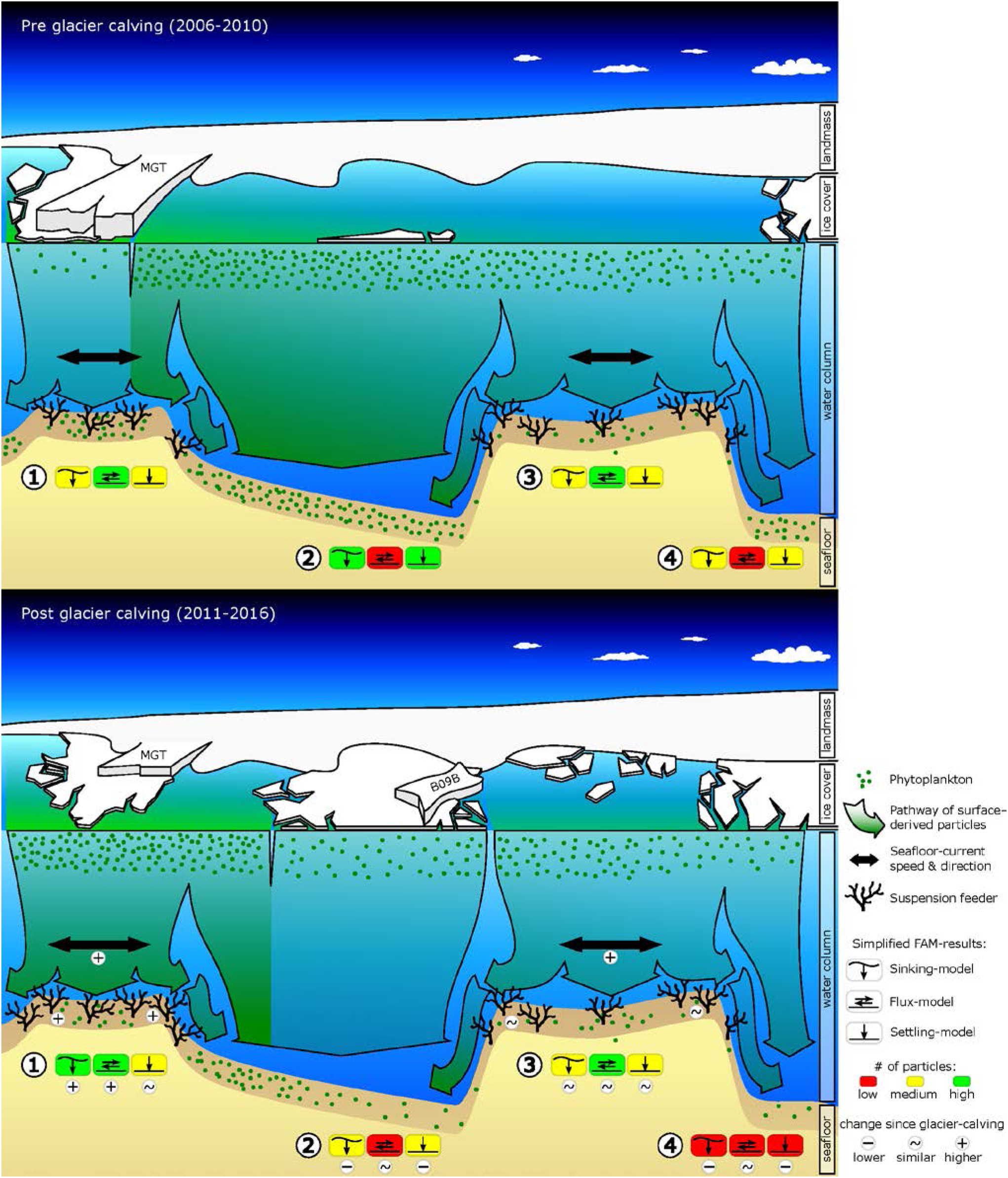
Graphic summarizing observed and predicted changes in environmental conditions (sea-ice, surface-chl-*a*, ocean current speeds, food-export) and seafloor fauna due to the calving of the Mertz Glacier Tongue (MGT) in 2010. The graphics shows a cross-section of the George V continental shelf approximately 80 km off the coast, looking South towards the Antarctic continent. The top graphic shows pre-calving environmental conditions and displays abundances of suspension feeders as observed from towed camera images. The bottom graphic shows observed changes in sea-ice, surface-chl-*a* and the position of the grounded iceberg B09B, as well as modelled changes in ocean current speeds, food-availability and suspension feeder abundances. Simplified food-availability-model (FAM) results are indicated for (**1)** Mertz Bank,(**2)** Adélie Sill,(**3)** Adélie Bank and (**4)** Adélie Basin. Additional indicators are included in the bottom graphic to highlight important changes

**Figure 2.**
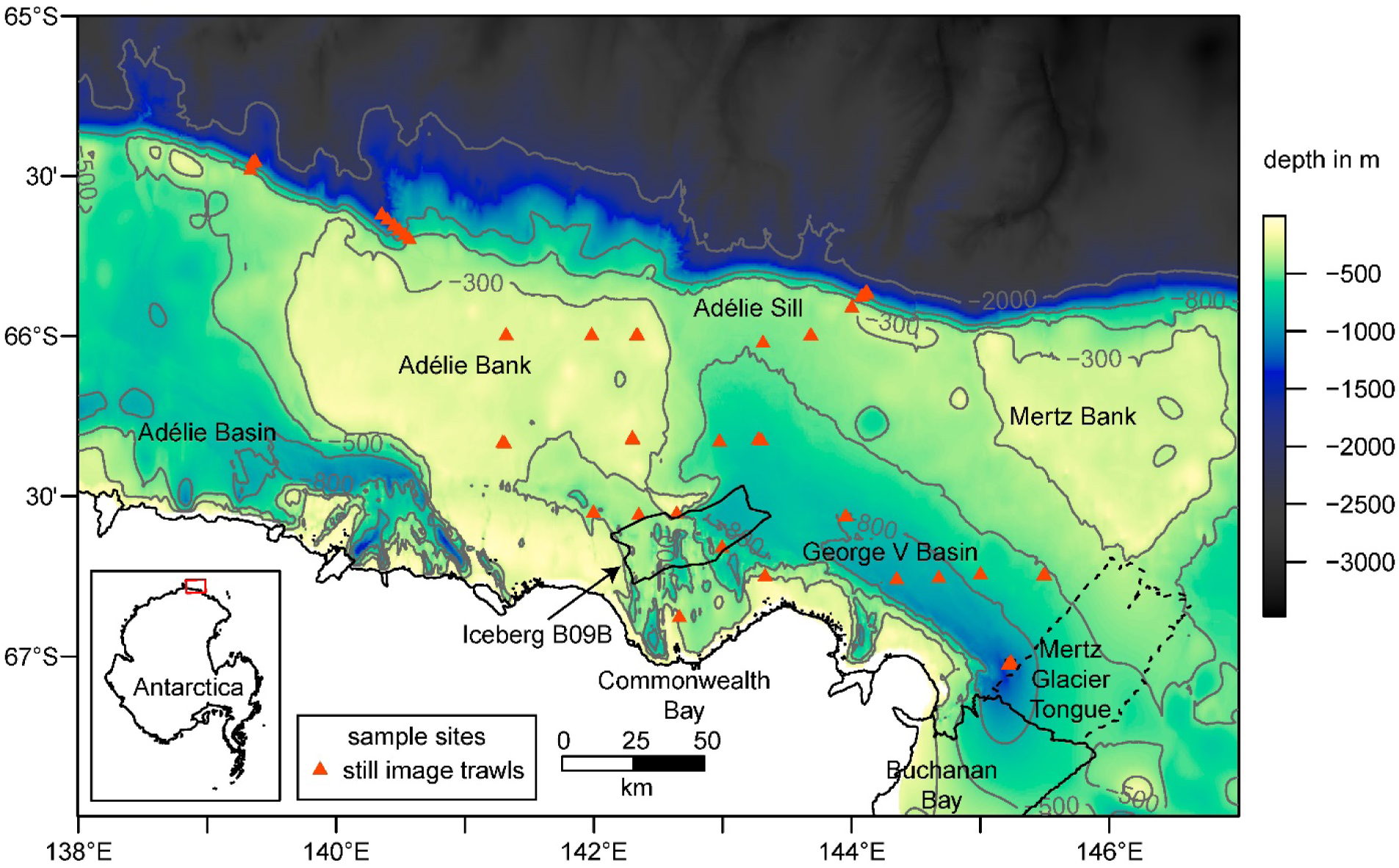
Bathymetry (Beaman et al., 2011) with selected contour lines (in grey), coastline and sample locations in the study area. Major glacial features such as the Iceberg B09B in the centre (location in 2016) and the Mertz Glacier Tongue (MGT) on the Eastern margin of the map are included. The dashed line shows the extent of the MGT prior to the calving in 2010. The inset map shows the location of the study area as highlighted by the red box.

## 2 Results

### 2.1 Changes in environmental conditions

Our results reveal that several aspects of the observed and the modelled marine environment have changed since the calving of the MGT(Fig. 3). Average sea-ice concentrations increased in the study region by 50-80%, except over the Mertz Bank and where the calved MGT was located (Supplementary Fig. 1). Near the Mertz Bank, average surface-chlorophyll-*a (*chl-*a*) concentrations increase by a factor of two or more (Fig. 3c), due to the change in availability and timing of ice-free surface waters that stimulate the growth of phytoplankton. The area of highest average surface-chl-*a* concentration also shows an eastward extension into areas previously covered by sea-ice. West of the Mertz Bank, the George V Basin and the Adélie Bank show decreasing values for surface-chl-*a (*Fig. 3c). In this area, the breakup of the sea-ice post-calving occurs much later in the year (Supplementary Fig. 1), shortening the time-period where surface phytoplankton is observed by satellites from around 4.5 months to 3 months. At the South-East tip of the Adélie Bank, north of the grounded giant iceberg B09B, the outline of a newly formed polynya (Tamura et al., 2012; Fogwill et al., 2016) can be observed, marked by lower sea-ice concentrations and higher surface-chl-*a*. Modelled seafloor current speeds (Supplementary Fig. 2) increase by about 5 cm/s on the shallower sections of the shelf down to around 500m depth, and decrease by almost 50 % at the shelf break and slope, as well as in the area previously occupied by the MGT. Changing bottom tidal current speeds account for almost all the increase in current speed on the South-Western flanks of the Adélie and Mertz Bank, in the deep George V Basin and below the iceberg B09B at the North-Western edge of Commonwealth Bay (Fig. 3f).

**Figure 3.**
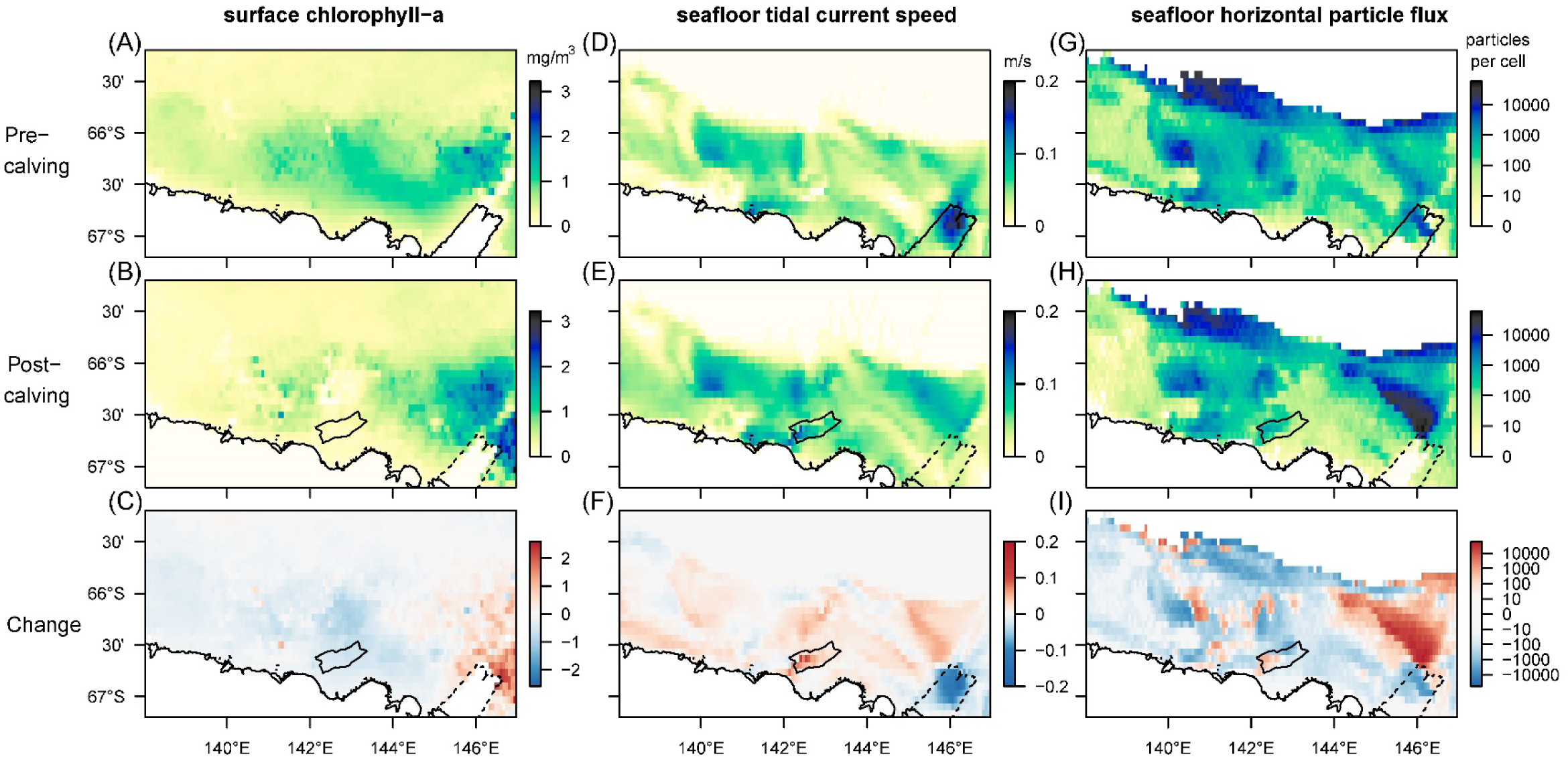
Comparison of mean values for selected biologically relevant environmental variables before and after the calving of the Mertz Glacier Tongue (MGT) in 2010. The first row shows environmental conditions in the five years leading up to the calving, the second row in the five years after, and the bottom row shows the magnitude of the change.(**A)/(B)/(C)** are satellite-derived (MODIS-A) estimates of surface-chl-*a* corrected for Southern Ocean application (Johnson et al., 2013). The missing data at the previous location of the MGT stems from a landmask-artefact in the NASA-data.(**D)/E/(F)** represent the speed of fluctuating currents (tidal currents) at the bottom layer of the regional ocean model used for this study (Cougnon et al., 2017).(**G)/(H)/(I)** show the number of particles moving horizontally along the seafloor before their permanent sedimentation (on log-scale). In all maps the strongest changes can be observed in the Eastern section of the region, close to the location of the MGT. In the post-calving maps, the outline of the newly grounded iceberg B09B is added for reference while dotted lines indicate the original position of the glacier tongue before it broke off.

The Food-Availability-Model (FAM) tracks and quantifies three components of surface-derived food particles: the sinking component captures the advection of phytodetrital matter by currents as it sinks through the water column until it reaches the seafloor; the flux component represents the horizontal flux of food particles along the seafloor before sedimentation; the settling component represents the final location of advected particles after taking into account the redistribution by seafloor currents. Sinking and settling particles follow similar patterns to the other environmental variables mentioned before, with an eastward shift for the peak number of sinking and settling particles, and absence of sedimentation on large parts of the Mertz Bank (closest to the former tip of the MGT) due to increased current speeds (Supplementary Fig. 3). Horizontal food flux along the seafloor, which is dependent mainly on the interaction between the distribution of surface productivity and seafloor current speeds, increases 20-50 fold on wide sections of the Mertz Bank (Fig. 3i). In contrast, changes are more patchy on the Adélie Bank, where increases in flux are mostly restricted to the inner section of the bank and the shelf break, while the edges of the bank experience decrease in flux. Further, most of the deeper sections of the shelf experience lower flux than before the calving.

### 2.2 Predicted changes in suspension feeder abundances

Mapped predictions of suspension feeder (SF) cover are based on the statistical relationship between pre-calving cover estimated from still-images, and the environmental covariates depth and log (horizontal flux)(Supplementary Table 1, deviance-explained = 44 %), which are selected as the best predictor variables by the stepwise regression process (Supplementary Table 2). There is a good fit between the predicted values from the statistical model and the observed values at the sampling sites, with a slight underestimation of high cover values (Fig. 4). SF-cover before the calving is high on most of the shallower sections of the shelf (<500 m depth)(Fig. 5a), with values estimated between 60-100 % cover. The two shallow banks in the study area, the Mertz Bank closest to the MGT in the East and Adélie Bank in the West, show relatively similar patterns of SF-cover, with average values of around 60 % except for on the edges of the banks, where cover of suspension feeders may reach up to 100 %. In contrast, post-calving predictions show a clear difference between the two banks (Fig. 5b) in that SF-cover is predicted to increase by 20-40 %(Fig. 5c) on large parts of the Mertz Bank, while the Adélie Bank and other areas retain similar SF-cover as previously. The stark difference in predicted SF-abundance between the Mertz Bank in the east and the Adélie Bank in the west stems from the 20-50-fold increase in particle flux on the Mertz Bank that is a direct result of both increased surface production and stronger tidal currents.

**Figure 4.**
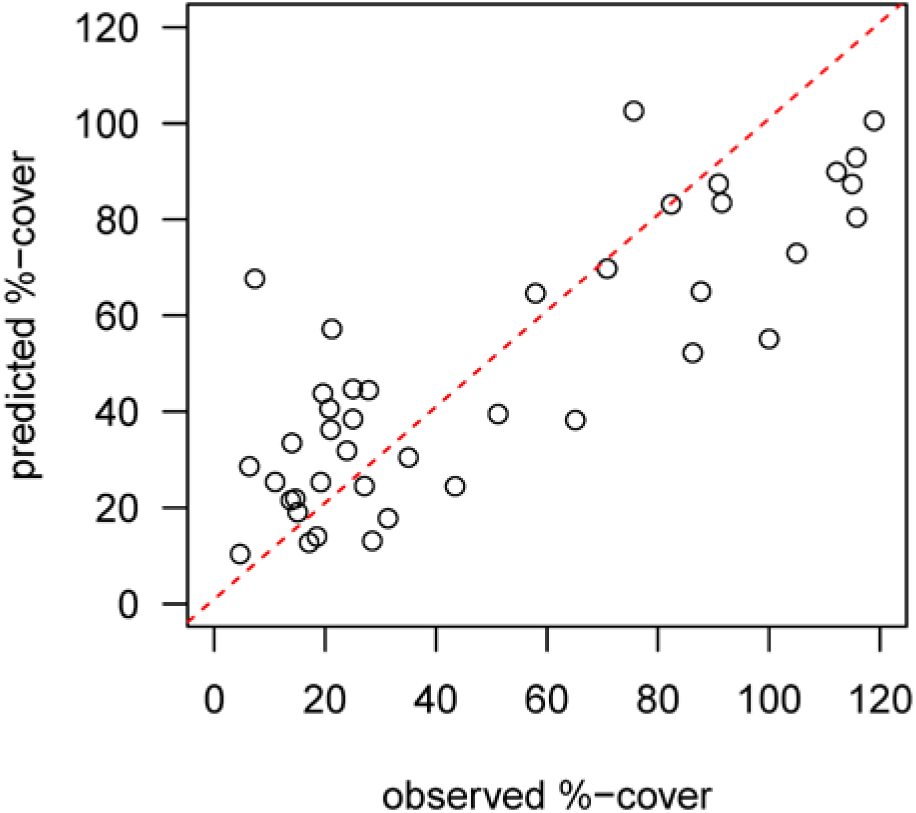
Diagnostic plot showing the fit between predicted values from the pre-calving statistical model and the observed %-cover estimates from benthic still images at the sample sites. The red dotted line indicates a perfect fit.

**Figure 5.**
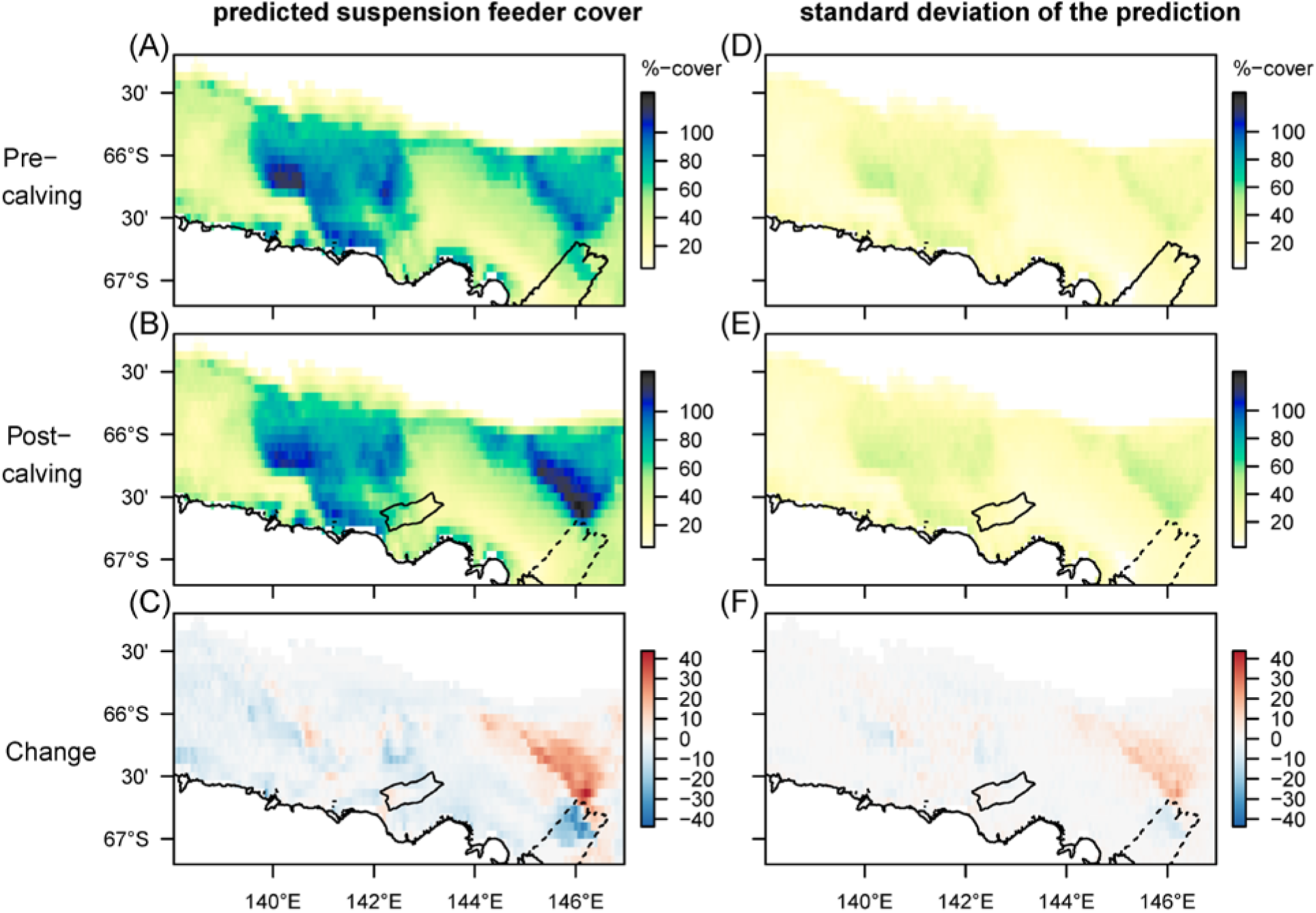
Spatial predictions of percentage cover of suspension feeders on the seafloor before and after the glacier calving. The cover of each species is estimated separately, thus overlapping cover can result in total coverage exceeding 100 %.(**A)/(B)/(C)** show the predicted mean, and (**D)/E/(F)** the standard deviation of the predictions. Note that changes are most pronounced on the Mertz bank close to where the glacier calving happened. The high predicted cover pre-calving under the glacier tip is driven by high current speeds, but samples have so far not been taken in this location.

## 3 Discussion

We predict that the calving of massive icebergs will have far-reaching effects on benthic communities mediated through the mechanism of pelagic-benthic coupling, and that changes occur even hundreds of kilometers away from the glacier tongue. While previous studies have shown that calving events can have localized negative impacts on the benthos through iceberg scouring (Gutt et al., 1996), here we predict that the combination of changes in local oceanography and surface production influences patterns of seafloor food-availability at much larger scales. Particularly strong changes are predicted in the horizontal flux of food particles post-calving, which is important in determining the distribution of suspension feeders (Jansen et al., 2018). Similar to other Antarctic regions that have recently become ice-free (e.g. Fillinger et al., 2013), environmental conditions on the Mertz Bank now seem much more favorable for suspension feeders (SFs) than before the calving. Our modelling suggests that there will be a strong, but locally confined increase in SF-abundance on the Mertz Bank of up to 40 %. Further away, near the Adélie Bank, increases in bottom current speeds seem to compensate for the overall decrease in food supply, resulting in a prediction of only marginal changes in SF-abundance. The distribution of surface production around the newly formed polynya on the leeward side of the grounded iceberg B09B(Fogwill et al., 2016) seems to slightly influence SF-abundances with relatively stable cover predicted beneath the polynya in contrast to decreasing cover in the ice-covered area directly north of the iceberg B09B. Close to and below the position of the tip of the MGT before it broke away, where Beaman and Harris (2005) have previously found a high number of macrobenthic species, including many sponges and bryozoans, our model predicts a substantial decrease in SF-abundance, due to a decrease in floor current speed affecting the horizontal food-flux. However, we caution that little confidence should be placed in this result; the environmental conditions in this area might be unique due to the glacier tongue and we lack biological samples for this area. Further, we also lack confidence in the food-availability-data because of missing data in the remotely sensed surface chl-*a* dataset (see Methods section 4.2.2). Unfortunately, the shallower sections of the Mertz Bank and the western part of the Adélie Bank, which we show here are particularly interesting areas, have not been physically sampled as part of the survey (see Fig. 2). However, because the survey was designed to cover a wide range of depths and geomorphologies (Hosie et al., 2011), because of the high confidence in the predicted values from the statistical model (Fig. 4), and because of the similarity in environmental conditions between the shallow banks prior to the glacier calving, we are confident the relationship between environmental variables and distributional patterns of suspension feeder abundances is consistent across the region.

Regions around ice-shelfs and glacier tongues provide valuable insight into the dynamic environment of the Antarctic shelf. When an ice-shelf calves a massive iceberg or collapses entirely, the marine environment, to which species might have acclimated to for many years, can transition quickly between a food-poor and a food-rich system (Gutt et al., 2011a). The MGT is thought to calve massive icebergs in a ∼70 year cycle (Giles, 2017), meaning that there are likely also differences in the long-term stability of environmental conditions, and in the frequency of iceberg-scour between the Mertz Bank near the MGT and the Adélie Bank in the West of the George V shelf. Studies on the West Antarctic Peninsula suggest that at least some components of Antarctic benthic communities on the shelf, such as glass sponges and pioneering species, can increase rapidly in areas that are newly ice-free, fueled by higher export of surface production (Gutt et al., 2011a; Fillinger et al., 2013). Conversely, slower-growing deep-sea corals and bryozoans may respond more slowly to changing environmental conditions. Whether there are any species or communities that have adapted to these long-term cyclic events in the George V region is unknown, because a comparative study between the Mertz and Adélie Bank has not been conducted. However, if the benthic community response in East Antarctica is similar to that of the West Antarctic Peninsula, the community composition on the Mertz Bank can be expected to change rapidly in the more favorable environment after the glacier calving, or will have undergone changes already, given that eight years have passed since the calving event. The postulated more favorable environment on the Mertz Bank might continue to persist for some time until the MGT regrows the ice tongue. Further, oceanographic models from after the calving indicate an increase in basal melting of the MGT due to warmer, faster moving waters from the east after grounded tabular iceberg relocation and the MGT calving (Cougnon et al., 2017) which may slow the regrowth of the MGT.

Food-availability is a key factor influencing species distributions. Here, we map predicted changes in relevant seafloor-food-availability caused by the calving of a major glacier tongue, and predict change in distributional patterns of benthic suspension feeders, a key element of Antarctic biodiversity. The study area on the George V shelf lies within the recently proposed East Antarctic MPA (AustralianAntarcticDivision, 2017), and we suggest the Mertz Bank and Adélie Bank should be considered as distinct areas for future sampling of the benthic community. The predicted distribution of suspension feeders after the glacier-calving provides an up to date picture of a key part of seafloor biodiversity, from which the representativeness of the proposed MPA can be assessed. Until regular monitoring programs are established, modelling studies such as ours give important information and context for future monitoring and assessment. Our study provides insight into temporal change and into the mechanisms that drive changes at the seafloor. This is important for a holistic understanding of the Antarctic marine ecosystem, and helps us to understand how climate change can affect the seafloor in the future.

## 4 Materials and Methods

### 4.1 Study area

The study area is located on the relatively deep (500-700 m) East Antarctic continental shelf and slope between latitudes 139°E and 147°E(Fig. 2). The depth of the shelf ranges between 200 m on the banks to 1300 m in the basins. The most prominent feature in this region is the Mertz Glacier Tongue (MGT) in the east which has strong influences on both oceanography (Barber and Massom, 2007) and biology (Arrigo and van Dijken, 2003; Sambrotto et al., 2003; Beans et al., 2008; Jansen et al., 2018). Strong katabatic winds in the region drive sea-ice production (Massom et al., 2001), convection of dense water that contributes to overturning circulation (e.g. Williams et al., 2008), and importantly also form ice-free surface areas on the westward sides of the MGT and the grounded icebergs. These permanent ice-free areas support a long growing season for phytoplankton resulting in high phytoplankton productivity (Arrigo and van Dijken, 2003; Sambrotto et al., 2003; Beans et al., 2008). Abundant and diverse benthic suspension feeder communities have been found primarily on the shallower section of the shelf between 200-600 m (Post et al., 2011). Further, tidal currents on the seafloor redistribute surface derived production, with flux rates of organic particles directly related to the abundance and species richness of the benthic community (Jansen et al., 2018).

The MGT calves off massive icebergs in an estimated 70-year cycle (Campagne et al., 2015), the last event happening in 2010 after a collision between the massive iceberg B09B and the MGT. Since the calving of the MGT, the Mertz Polynya has decreased significantly in size (Tamura et al., 2016), changing ocean circulation (Aoki et al., 2017; Cougnon et al., 2017; Kusahara et al., 2017) and increasing sea-ice concentrations (Tamura et al., 2012). The iceberg B09B grounded on the South-Eastern flank of the Adélie Bank shortly after the collision, and a new polynya has formed on its leeward side (Fogwill et al., 2016). For more details on the study area and the oceanography we refer to numerous papers on the region (e.g.Beaman and Harris, 2005; Cougnon et al., 2013; Shadwick et al., 2013; Cougnon et al., 2017; Kusahara et al., 2017; Jansen et al., 2018).

### 4.2 Environmental data and numerical modelling

#### 4.2.1 Ocean model and bathymetry

Ocean current speeds and directions before and after the glacier calving are derived from a tide-simulating oceanographic model for the George V shelf developed by Cougnon et al.(2017) based on the Regional Ocean Modelling System (ROMS)(Shchepetkin and McWilliams, 2005). The model setup used here is similar to that described by Cougnon et al.(2013), using the same horizontal and vertical grid. The horizontal grid has a resolution of 2.16 km near the southern boundary and 2.88 km near the northern boundary. The vertical grid is arranged to give higher resolution at the top and bottom of the water column. The model domain encompassed the area from the Antarctic coastline to the deep ocean at 62.72°S, and from 135.77°E to the west of the French base, Dumont D’Urville, to 158.08°E to the east of George V Land (Cougnon et al., 2013).

The model includes ocean-ice shelf thermodynamics described by three equations following Holland and Jenkins (1999), frazil ice thermodynamics following Galton-Fenzi et al.(2012), as used in previous studies (e.g.Cougnon et al., 2013; Gwyther et al., 2014) and a simplified analytic tidal forcing at the lateral boundaries (Cougnon et al., 2017 for details). The bathymetry in both simulations is based on RTopo-1(Timmermann et al., 2010), and modified to include local high-resolution bathymetry (Beaman et al., 2011) as described in Mayet et al.(2013). Ice draft of the MGT and B09B, along with the underlying bathymetry, is based on an early version of the most up-to-date product by Mayet et al.(2013).

The total run time of the model for each simulation (before and after calving) is 33 years, using an annually repeating loop of the same lateral forcing for both simulations and an annually repeating loop of the surface forcing corresponding to each icescape. This 33 year run includes a spinup phase of 30 years to reach equilibrium.

#### 4.2.2 Surface productivity and sea-ice

We estimated spatial patterns of surface productivity from measures of ocean colour derived from NASA’s Moderate Resolution Imaging Spectroradiometer (MODIS-Aqua)(NASA Goddard Space Flight Center, 2014). We used Level-3 binned daily remote sensing reflectance, provided at a resolution of 4km equal-area bins, and corrected the values for Southern Ocean application using the algorithm in Johnson et al.(2013). Daily measures of chlorophyll-a concentrations were averaged for southern hemisphere spring and summer in each year for a five-year period before (2005-2009) and after (2011-2016) the calving of the MGT. There are no data available from NASA for the area previously covered by the MGT, presumably due to a landmask artefact.

We estimated seasonal patterns of sea-ice concentrations from satellite-measures of Nimbus-7 SMMR and DMSP SSM/I-SSMIS Passive Microwave Data (Cavalieri et al., 1996, updated yearly). Daily measures of sea-ice concentrations were averaged for southern hemisphere spring and summer in each year for a five-year period both before (2005-2009) and after (2011-2016) the calving of the MGT.

#### 4.2.3 Food availability model (FAM)

We mapped the availability of surface-derived food at the seafloor before and after the calving event using a validated food-availability model (FAM) as described in Jansen et al.(2018). The FAM uses a distribution of particles that is based on multi-year averages of satellite derived chlorophyll-a (section 4.2.2), and tracks individual particles from the surface to the seafloor while accounting for their sinking speed, the speed and direction of currents (section 4.2.1) in 3D, and the sedimentation-rate of particles on the seafloor based on particle sizes (Jansen et al., 2018). The model generates three maps of food availability, namely a sinking-map (showing the number of particles arriving/temporarily settling on the seafloor), a map of horizontal flux (showing where particles move along the seafloor before their sedimentation), and a settling-map (showing where particles permanently settle on the seafloor). The only difference from our approach here to the Jansen et al.(2018) model is the use of a different ocean model and a different surface-chl-*a* climatology. Current speeds on the shelf seem to be slightly lower in the pre-calving part of the model developed by Cougnon et al.(2017) compared to that of Cougnon et al.(2013), but are within the seasonal variation of the Cougnon et al.(2013) model.

For the particle tracking, we used four consecutive time-slices of the ocean-model for the summer season before and after the calving respectively. We used the four consecutive time slices with the strongest differences in current direction and speed, to ensure that each time slice adequately captures one full tidal movement (a 6h time-slice with 3 hours of incoming tide and 3 hours of outgoing tide would show very little current speed). The maximum number of seed particles was ∼4.5 million for the pre-calving model and the particles were tracked in 30 min time-steps. At each time-step the location of each particle with respect to the ROMS-cells was calculated, and water current speed and direction at that location updates for advection of the particles during the next time-step. Particles were stopped when they either moved out of the study area or matched the stopping criteria for the respective model, as described in Jansen et al.(2018). The resulting particle distributions from each model-run were back-transformed into a regular grid with a resolution of 1/15 degrees.

We use the FAM-parameters previously defined for this region (Jansen et al., 2018), namely a sinking speed of 300 m/day, a particle radius of 0.24 mm, the density of seawater at 1030 kg/m^3^, the density of settling particles at 1100 kg/m^3^ and an aspect ratio of 1 representing idealised spherical particles in our modelling.

For the particle tracking, we used R Version 3.3.1(R Core Team, 2016) with the packages “ptrackr” (Jansen and Sumner, 2017), “raster” (Hijmans and 2015), “ncdf4” (Pierce and 2014), “nabor” (Elseberg et al., 2012), “geostatspat” (Brown, 2015) and “spatstat” (Baddeley and Turner, 2005).

### 4.3 Biological data collection

We use the same dataset of benthic images as used by Jansen et al.(2018), which is available through the Australian Antarctic Division Data Centre (Jansen et al., 2017). It comprises detailed underwater still images collected during the Collaborative East Antarctic Marine Census (CEAMARC) for the Census of Antarctic Marine Life in December 2007 to February 2008(Hosie et al., 2011). Transects during the CEAMARC were designed to cover a wide range of depths and geomorphologies in the region and therefore can be considered representative of the area modelled. A forward facing 8 megapixel Canon EOS 20D SLR with two speedlight strobes was mounted on a beam trawl and pictures were taken every 10 seconds. 32 sites were sampled with transect length mostly between 4-6 km, with exceptions ranging between 3-16 km. The trawl was controlled using a deck winch. Benthic fauna were identified to the lowest taxonomic resolution possible and, where species identification was not possible, specimens with similar overall appearance were grouped into morphotypes. The bottom third of each image was scored. For each image, the abundance of each species/morphotype was estimated within 5% bins from 0% to 50% and 10% bins from 50% to 100%. Using taxonomy and body-type along with expert knowledge, the abundance of the suspension feeding fauna in each picture was calculated.

### 4.4 Statistical Analysis

We use the image data and the maps of environmental data for 2005-2009 to generate a pre-calving statistical model, aiming at producing the statistical model that best explains the abundance of suspension feeders (SF). Each transect was split at the boundaries of the environmental grid cells to ensure all pictures lay within the same value for the environmental covariates. We multiplied %-cover estimates in each image by 100 and rounded up to generate integer values that better suit a statistical analysis using a multiple linear regression with a negative binomial GLM(assuming the values would then represent the number of pixels covered by the fauna). We backwards selected variables from a full model using AIC. The full model contained the important environmental variables identified by Jansen et al.(2018), namely depth, tidal-current speed and the horizontal flux of particles along the seafloor. The final model contained only depth and log (horizontal-flux) as predictor variables. We found that using a negative binomial generalized linear model (compared to a linear model in the previous study) did not affect the selection of model terms. Therefore, the change in selected model terms is likely to come from the difference in the ocean model or the surface productivity. The pre-calving statistical model showed a good fit between the predicted and the observed values at the sample sites, with possibly a slight underestimation of suspension feeders at high abundances (Fig. 5).

We then used the pre-calving statistical model to predict the spatial distribution of SF-cover in both the pre-calving and the post-calving environment. The difference between the resulting maps was used to make inferences about areas with expected increases and decreases in the abundance of suspension feeders. Further, we bootstrapped the parameters of the pre-calving statistical model to obtain estimates for the standard-deviation of the predictions.

For the statistical analysis, we used R Version 3.3.1(R Core Team, 2016) and the packages “raster” (Hijmans and 2015), “MASS” (Venables and Ripley, 2002), “maptools” and “modEvA”.

### 4.5 Data availability

Estimates of suspension feeder abundances from benthic images are available through the Australian Antarctic Division Data Centre (Jansen et al., 2017). Raster files containing mapped predictions of food-availability and suspension feeder abundances presented in this study, from before and after the glacier calving are available through the Australian Antarctic Data Centre (Jansen, 2018).

## 5 Acknowledgements

We thank Marc Eléaume for useful discussions and for making the original biological data available to us. Biological samples were collected during the CEAMARC program as part of the IPY #53 Census of Antarctic Marine Life program. Coastline and glacial features for the figures are taken from the Antarctic Digital Database version 5. J.J. is supported by a Tasmanian Graduate Research Scholarship and a QAS Top-Up scholarship. This work was completed as part of Australian Antarctic Science project 4124.

## 6 Author contributions

Concept/design of study: JJ, NH, CJ, PD, EC Analysis of data: JJ, NH, CJ, PD

Numerical modelling: EC, BG-F Figures and tables: JJ

Writing paper: JJ with contributions from all authors

